# Germline loss diminishes somatic mitochondria but confers preservation of respiratory function during aging and hypothermia

**DOI:** 10.1101/2024.09.20.614121

**Authors:** HyeJin Hwang, Brandon J Berry, Claudette St. Croix, Andrew P Wojtovich, Sruti Shiva, Arjumand Ghazi

## Abstract

Reproductive status influences metabolism and health across lifespan in diverse ways and mitochondrial function playing a critical role in mediating this relationship. Using the *Caenorhabditis elegans* germline ablation model, we investigated the impact of germline stem cell (GSC) loss on mitochondrial dynamics and respiratory function. Our results show that GSC loss reduces mitochondrial volume and respiratory function in young adulthood but preserves mitochondrial activity during aging and upon exposure to hypothermic stress, correlating with enhanced survival. We found that the transcription factor NHR-49/PPARα, but not DAF-16/FOXO3A, was essential for preserving mitochondrial function and hypothermia resistance in these long-lived mutants. Together, these findings reveal the impact of germline signals on somatic mitochondrial health and underscore the intricate relationship between reproductive fitness and organismal health.

## INTRODUCTION

Reproduction and aging are linked through a complex and dynamic relationship shaped, in part, by metabolism and energetics. Reproductive status influences whole-body metabolism across the lifespan in women as puberty, childbirth, reproductive dysfunction and menopause alter lipid homeostasis and energy balance and shape overall health. Menopause in women is often accompanied with fat reorganization, weight gain, metabolic slowdown, and an increased risk of age-related diseases such as diabetes and cardiovascular disease CVD) (El Khoudary, 2020). Obesity and low body fat contribute to ∼12% of female infertility, while conditions like endometriosis and polycystic ovarian syndrome (PCOS) trigger systemic metabolic dysfunction with severe disease consequences (Capezzuoli et al., 2022). Interestingly, PCOS and primary ovarian insufficiency (POI) are sometimes linked to reduced risks of hormonal cancers but also increased muscle mass (Harris et al., 2019; Kogure et al., 2015). While early centenarian studies suggested that childlessness correlated with increased longevity (Westendorp and Kirkwood, 1998), about half of female centenarians have been noted to give birth in their forties (Perls et al., 1997), suggesting that late-life fertility might signal slower aging. Research in laboratory animals supports the complex reproduction-longevity relationship too. Sterility is often correlated with longevity and mating reduces lifespan and stress resistance in many species (Fowler and Partridge, 1989; Maures et al., 2014; Shi and Murphy, 2014; Zwaan et al., 1995). Yet, genetic studies in worms and flies, and ovarian transplantation in mice, have revealed the benefits of reproductive signals on stress resilience, longevity, and age-related fitness (Amrit and Ghazi, 2017; Flatt et al., 2008; Hsin and Kenyon, 1999; Mason et al., 2009; Mason et al., 2022). The mechanisms by which metabolism influences, and is shaped by, the fertility-longevity-health relationship remain poorly understood.

Mitochondria play crucial roles in mediating the link between reproduction and longevity as they regulate energy production, metabolic homeostasis and signaling, which are essential for reproductive fitness, stress response and healthy longevity (Miwa et al., 2022). There is compelling evidence that signals from somatic tissues influence mitochondrial status in germ cells to influence reproductive output and organismal health. Nutrient sensing pathways such as insulin/IGF1 signaling and target of rapamycin (TOR) alter mitochondrial dynamics within germ cells and thus impact fertility in multiple species (Das and Arur, 2017; Lopez et al., 2013; Sieber et al., 2016; Templeman and Murphy, 2018). Disruptions of somatic mitochondrial homeostasis, such as through prolonged activation of the mitochondrial unfolded protein stress response pathway (UPRmt) or inactivation of mitochondrial electron transport chain (ETC) activities, influence germline health and fertility (Charmpilas et al., 2024; Rea et al., 2007; Xu et al., 2018). Recent studies have suggested that pro-longevity perturbations, including those involving germline loss, trigger changes in mitochondrial dynamics, and germline mitochondria have been reported to play a role in coordinating an organism-wide UPRmt response to acute mitochondrial stress (Chaudhari and Kipreos, 2017; Palikaras et al., 2015; Shen et al., 2024; Weir et al., 2017). Modulation of somatic mitochondria by reproductive signals likely impacts efficiency of energy utilization, cellular integrity and organismal fitness. But how reproductive signals impact mitochondrial energy utilization and how this shapes cellular integrity and organismal fitness across the lifespan remain unexplored.

In the nematode *Caenorhabditis elegans*, removal of germline stem cells (GSCs), which produce sperm and oocytes, leads to a significant extension of lifespan and enhanced stress resistance (Amrit and Ghazi, 2017; Arantes-Oliveira et al., 2002; Ghazi, 2013; Hsin and Kenyon, 1999). Similar life extensions have been observed in various species including flies, fish, and mice, suggesting a conserved biological mechanism (Flatt *et al*., 2008; Mason *et al*., 2009; Mason *et al*., 2022; Moses et al., 2024). The longevity of germline-ablated *C. elegans* is not merely due to sterility but is specifically linked to the elimination of GSCs, which normally emit signals that synchronize reproductive status with organismal health (Amrit and Ghazi, 2017; Arantes-Oliveira *et al*., 2002; Ghazi, 2013). Upon GSC removal, the organism undergoes systemic metabolic reprogramming that converts the reproductive challenge into an advantageous lifespan and stress-resistance enhancement. This adaptive response includes elevation of cytoprotective processes such as proteasomal function, autophagic flux and multiple stress-response pathways (Amrit and Ghazi, 2017; Ghazi, 2013). Lipid metabolism plays a pivotal role in this transformation, as several studies, including ours, have shown that lipid homeostasis is significantly altered following GSC removal (Amrit et al., 2016; Amrit and Ghazi, 2017; Antebi, 2013; Goudeau et al., 2011; Steinbaugh et al., 2015). Upon germline loss, a transcriptional network is activated in somatic cells that orchestrates the upregulation of lipogenic- and lipolytic-gene expression to mediate longevity and stress resistance (Amrit and Ghazi, 2017; Antebi, 2013; Ghazi, 2013). This network involves transcription factors such as DAF-16/FOXO3A, essential for diverse pro-longevity interventions including germline loss, as well as stress-response mediators, ligand-activated nuclear hormone receptors (NHRs) and a transcription elongation and splicing factor, TCER-1/TCERG1 (Amrit and Ghazi, 2017; Antebi, 2013; Ghazi et al., 2009). Previously, we identified the role of NHR-49, which performs functions undertaken in vertebrates by peroxisome proliferator-activated receptor alpha (PPARα). NHR-49 promoted germline-less longevity by upregulating the expression of genes involved in mitochondrial- and peroxisomal-β-oxidation and fatty-acid desaturation (Ratnappan et al., 2014; Ratnappan et al., 2016). Other pro-longevity factors such as DAF-16/FOXO, TCER-1/TCERG1 and SKN-1/NRF2, also mediate the upregulation of both lipogenic and lipolytic transcriptional profiles (Amrit *et al*., 2016; Amrit and Ghazi, 2017; Goudeau *et al*., 2011; Ratnappan *et al*., 2014; Steinbaugh *et al*., 2015).

Our studies suggested that, through a concerted enhancement of lipid anabolism and catabolism, NHR-49 and other factors mediate break down of fats allocated for reproduction, restoring lipid homeostasis and enhancing stress resistance and longevity. How these ostensibly antagonistic gene-expression changes translate into metabolic outcomes remains unclear. Here, we examined the impact of GSC loss on mitochondrial morphology and respiratory function and the role of NHR-49 and DAF-16 in these changes. We found that GSC-elimination results in reduction of mitochondrial volume and respiratory function. However, unlike normal fertile animals, GSC-less animals retained mitochondrial function with age and upon experiencing hypothermic stress, dependent largely upon NHR-49/PPARα but not DAF-16/FOXO. Overall, our studies underscore the complex relationship between mitochondrial dynamics, function and short- and long-term physiological outputs at the organismal level.

## RESULTS

### Germline loss causes reduction in mitochondrial volume and activity

We compared mitochondrial content and function between wild-type animals (WT) and temperature-sensitive mutants of *glp-1*, a gene essential for germline proliferation. *glp-1* mutants lose all their germ cells when grown at the non-permissive temperature (25oC) and are routinely used as a model for germline-less longevity (Arantes-Oliveira *et al*., 2002). We focused on mitochondria in the worm body wall muscles since skeletal tissue is a respiration intensive tissue whose diminished function is closely linked with aging (Xu and Wen, 2023). To assess mitochondrial organization and content, we used a well-established reporter expressing mitochondria-targeted GFP in body-wall muscles, *Pmyo-3::mtGFP* (Benedetti et al., 2006). Surprisingly, confocal image analysis and quantification of this reporter in single muscle cells showed that mitochondrial volume of young Day 1 *glp-1* adults was almost half that of muscle-cell volume of age-matched WT animals (Fig. 1A). In young, healthy animals, mitochondria are arranged in well-ordered, highly interconnected networks. However, the mitochondria of Day 1 *glp-1* mutants appeared to be arranged in dispersed clusters rather than orderly networks (Fig. 1B, top panels) although sphericity, another measure of mitochondrial morphology that takes surface area and volume into account, did not show a statistically significant difference between the two strains (Fig. S1A). Increased mitochondrial connectivity is often associated with improved mitochondrial function (Weir *et al*., 2017), so these observations suggested that the long-lived mutants’ mitochondrial function may be reduced as well. To test this, we measured whole-animal oxygen consumption rates (OCR) in live animals of both strains using SeaHorse technology (Yoo et al., 2024). In Day 1 *glp-1* mutants, basal respiration (BR), which reflects mitochondrial activity under normal conditions, was indeed lower than in their age-matched WT counterparts (Fig. 1C). The mutants also showed lower spare respiratory capacity (SRC)- the ability to synthesize ATP by oxidative phosphorylation in response to increased energy demand (Fig. 1D). In an alternative approach, we isolated mitochondria from Day 1 adults of these strains and measured mitochondrial oxygen consumption using a Clark-type Oxygen electrode (Berry et al., 2023; Clark et al., 1953). Mitochondria of *glp-1* mutants showed lower BR and SRC than WT in this assay as well (Fig. S2).

**Figure 1.**
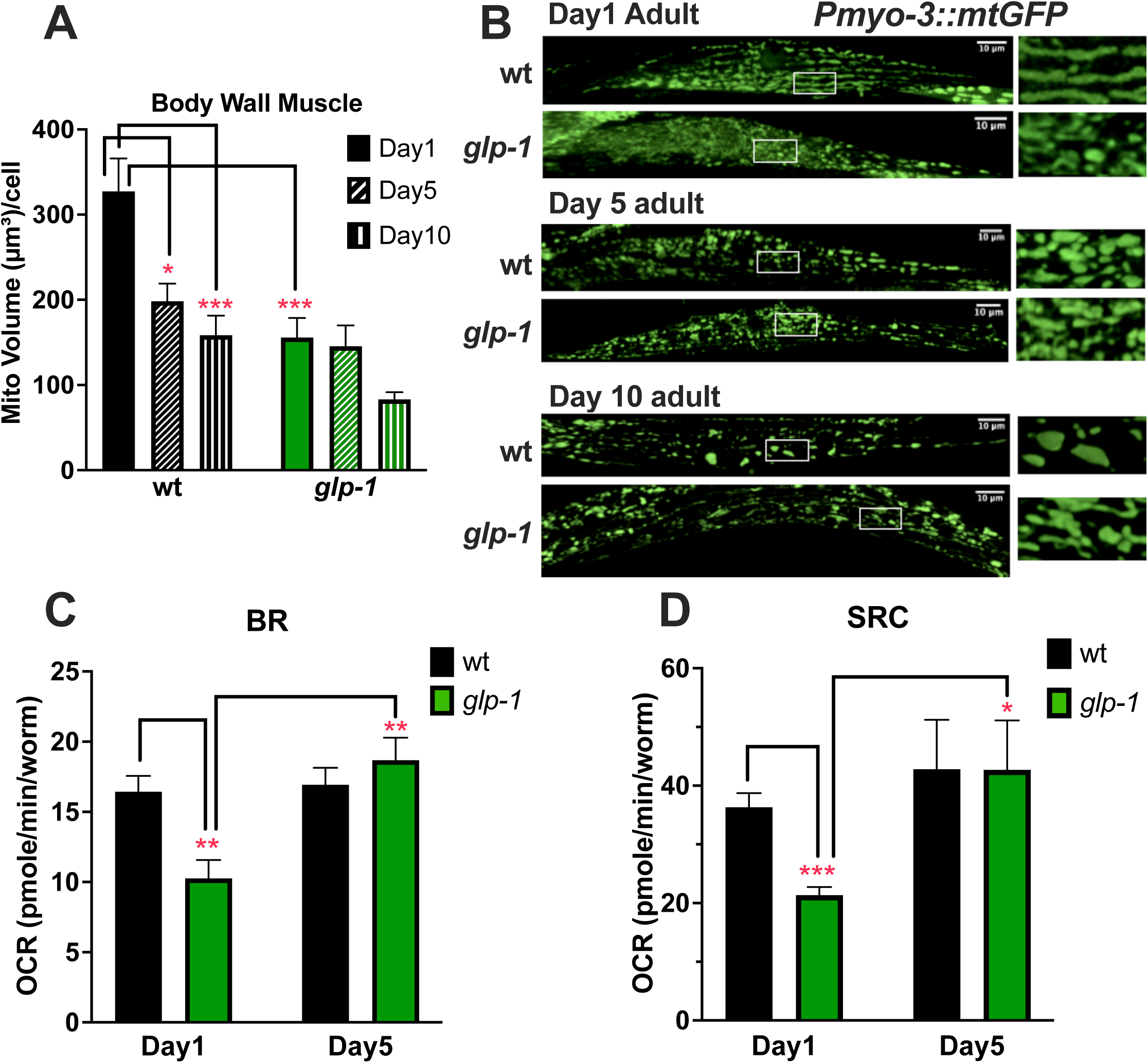
Germline-less animals have reduced mitochondrial volume and activity in young adulthood but maintain respiratory function with age. **A-B.** Mitochondrial volume is reduced and mitochondria appear fragmented in young *glp-1* mutants. Confocal images of the body wall muscle cell expressing Pmyo-3::mtGFP signals in wild-type (WT, black bars) and *glp-1* mutants (green bars) on Days 1(**B**, top), 5 (**B**, middle) and 10 (**B**, bottom) adults. Left panels show a single body wall muscle cell and white box indicates the area enlarged in the corresponding insets on the right. Scale bar indicates 10 µm. Quantification of this data is shown in A. Mitochondrial volume shown as average mean ± standard errors for one cell per animal. Total number of cells/animals examined in multiple independent experiments: (Day1 WT: n=16; Day 5 WT: n=17; Day10 WT: n=19; Day1 *glp-1*: n=16; Day5 *glp-1*: n=15; Day10 *glp-1*: n=12) Statistical significance calculated by two-way ANOVA followed by Tukey’s multiple comparison test. C, D. Mitochondrial respiration is reduced in Day 1 *glp-1* mutants but maintained with age. Oxygen consumption rate (OCR) measured in Day 1 and Day 5 adults using a Seahorsebio XF96 analyzer. Mitochondrial basal respiration (BR, C) and spare respiratory capacity (SRC, D) are indicated as OCR in average mean ± standard errors (number of trials: Day 1: n=5, Day 5: n=4). Statistical significance calculated by Student’s T-test and indicated by asterisks as *p<0.05, **p<0.01 and ***p<0.001.

### Germline-less, long-lived mutants maintain mitochondrial volume and activity with age

The well-ordered networks and connectivity of mitochondria seen in young animals is progressively lost and their volume is reduced with age (Fig. 1A and 1B, top panels) (Weir *et al*., 2017). We compared the rate of this decline between the normal and long-lived strains. Expectedly, WT animals underwent ∼40% reduction in mitochondrial volume by Day 5 and then a further significant diminution by Day 10 (Fig. 1A). Morphologically, these changes were visible as reduced prevalence of tubular mitochondria and increase in spherical structures (Fig. 1B, middle and bottom panels). Strikingly, the *glp-1* mutants showed no reduction in mitochondrial volume by Day 5 and a small one by Day 10 that did not achieve statistical significance (Fig. 1A). Interestingly, mitochondrial DNA (mtDNA) copy number did not change significantly with age in wild-type or *glp-1* mutants, though *glp-1* mutants showed an expectedly lower content that WT due to loss of mitochondria-dense germ cells (Fig.S1B). Upon assessing the rates of respiration as the animals aged, we found that while there was no measurable reduction in WT animals’ BR, *glp-1* mutants’ BR was in fact modestly elevated with age such that by Day 5 the two strains manifested similar BR rates (Fig. 1C). Similarly, the SRC of *glp-1* mutants was also increased by Day 5 (Fig. 1D).

Together, these observations suggested that at the onset of adulthood, germline-less mutants exhibited reduced mitochondrial quantity and functionality. But, unlike WT animals, they maintained their mitochondrial status with age and even underwent a modest improvement.

### Germline-less, long-lived animals preserve mitochondrial function, and survive longer, upon exposure to hypothermia

Germline loss not only increases lifespan but also enhances resistance against multiple stressors (Amrit and Ghazi, 2017; Antebi, 2013; Ghazi, 2013). The preservation, and even enhancement, of mitochondrial function with age exhibited by *glp-1* mutants led us to ask if they also maintained respiratory capacity when subjected to environmental stress. Since SRC represents the ability to produce extra ATP under situations of sudden energy demands, we focused on hypothermia, a precipitous cold exposure during which cellular activities and metabolic rates are reduced in poikilothermic animals such as *C. elegans*. We exposed Day 1 WT and *glp-1* adults to 4oC for 4h and measured respiration rates. In WT animals, BR did not decline significantly upon cold exposure whereas SRC underwent ∼34% reduction (Fig. 2A, B). However, in *glp-1* mutants this SRC decline was not observed and both BR and SRC were preserved after at least 4h of cold stress in Day 1 (Fig. 2A, B) and Day 5 (Fig. 2C, D) *glp-1* mutants. This enhanced mitochondrial function was also accompanied with improved hypothermic survival. *glp-1* mutants survived significantly longer than WT when subjected to acute (4h) or prolonged (24h or 48h) exposure to 4oC (Fig. 2E, Table S1). Indeed, while at 20^0^C, *glp-1* mutants showed a ∼50% lifespan extension, after 48h of hypothermia, they survive ∼300% longer than wild type. Thus, germline loss resulted in lower mitochondrial volume and function during young adulthood under normal environmental conditions, but these modalities were preserved during aging and hypothermic stress, and it conferred the ability to maintain respiratory capacity and survive longer under stressful conditions.

**Figure 2.**
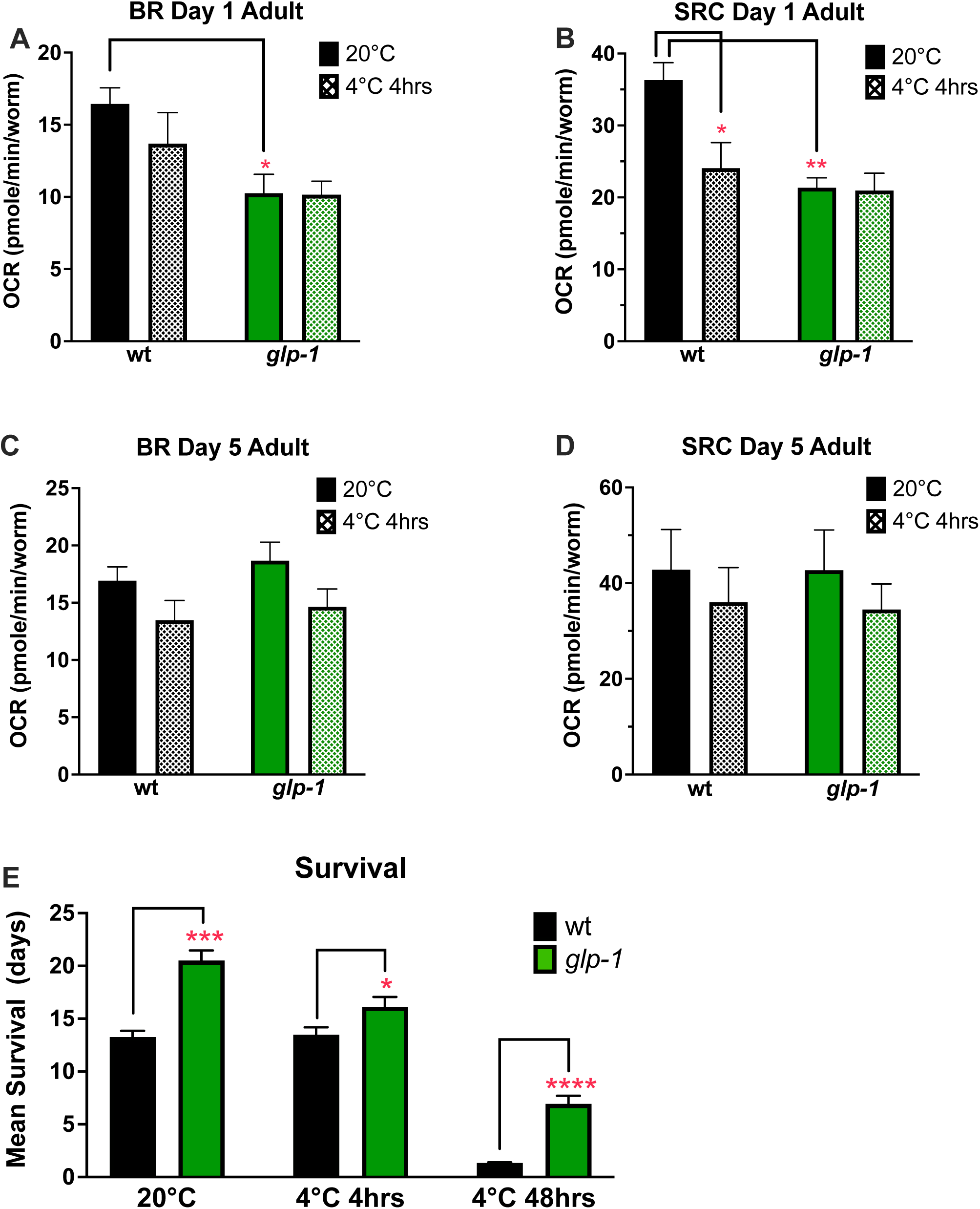
Germline-less animals retain mitochondrial function with age and survive longer upon hypothermic exposure. **A-D: *glp-1* mutants exhibit reduced oxygen consumption rate (OCR) but maintain it over time.** OCR measured using a Seahorsebio XF96 analyzer in Day1 (A, C) and Day 5 (B< D) adults of wild type (WT, black bars) and *glp-1* mutants (green bars) at normal temperature (20^0^C, solid bards) and upon exposure to cold shock (4oC, 4h, hashed bars). Mitochondrial basal respiration (BR, **A** and **C**) and spare respiratory capacity (SRC, **B** and **D**) are indicated as OCR in average mean ± standard errors (number of trials: Day 1=5, Day 5 =4). Data indicated was aggregated from 5 (Day 1) or 4 (Day 5) independent trials and statistical significance calculated by two-way ANOVA followed by Tukey’s multiple comparison. **E. *glp-1* mutants exhibit enhanced survival upon cold-shock exposure.** Survival of the same strains at 20^0^C and at 4^0^C after 4 hours indicated on Y axis. X axis shows age of animals. Analyses performed using Kaplan Meier survival statistics. Statistical signicfiance estimated by Mantel Cox test and shown as * p<0.05, ** p<0.01, ***p<0.001 and ****: p<0.0001. Representative data from one of three trials. Data from additional trials in Table S1.

### NHR-49 promotes the preservation of mitochondrial function in aging germline-less animals

Previously, we identified *nhr-49* as being essential for the longevity of germline-less animals (Ratnappan *et al*., 2014; Ratnappan *et al*., 2016). Since our studies in *glp-1* mutants, and other investigations, showed that NHR-49 mediated the upregulation of genes involved in mitochondrial fatty-acid β-oxidation (Ratnappan *et al*., 2014; Van Gilst et al., 2005a; Van Gilst et al., 2005b), we asked if it contributed to the maintenance of mitochondrial function with age and the enhanced survival of *glp-1* mutants upon hypothermia. In Day 1 adults, *nhr-49* mutation caused a significant reduction in the BR of WT animals but did not impact BR of *glp-1* mutants significantly. However, by Day 5, while *glp-1* mutants exhibited a perceptible increase in BR, *nhr-49;glp-1* showed a significant reduction (Fig. 3A). Similarly, the SRC increased manifested by *glp-1* mutants between Day 1 to Day 5 was abrogated in *nhr-49;glp-1* mutants (Fig. 3B). We could not measure the SRC of Day 5 *nhr-49* single mutants as they died rapidly upon exposure to the mitochondrial uncoupler DCCP used in this assay. We next assessed the role of NHR-49 in germline-less animals’ ability to maintain respiratory capacity under hypothermia. Expectedly, while *glp-1* mutants showed lower BR and SRC than wild type at 20^0^C, these functions were maintained upon exposure to 4^0^C for 4h, in contrast to the significant decline undergone by WT animals (Fig. 3C, D). *nhr-49* single mutants underwent a much greater decrease in SRC than WT animals upon cold stress (34% reduction in WT and 67% reduction in *nhr-49* mutants compared to same strains at 20°C) (Figure 3D). SRC of *nhr-49;glp-1* though lower was not statistically different from *glp-1*. Together, these data suggested a specific impact of NHR-49 on respiratory function, based on germline status and age of the animal as well as the stress being experienced. In normal, fertile animals, it was critical for basal respiration at all ages and under hypothermia. But, in long-lived mutants, it was specifically required for maintenance of mitochondrial function with age.

**Figure 3.**
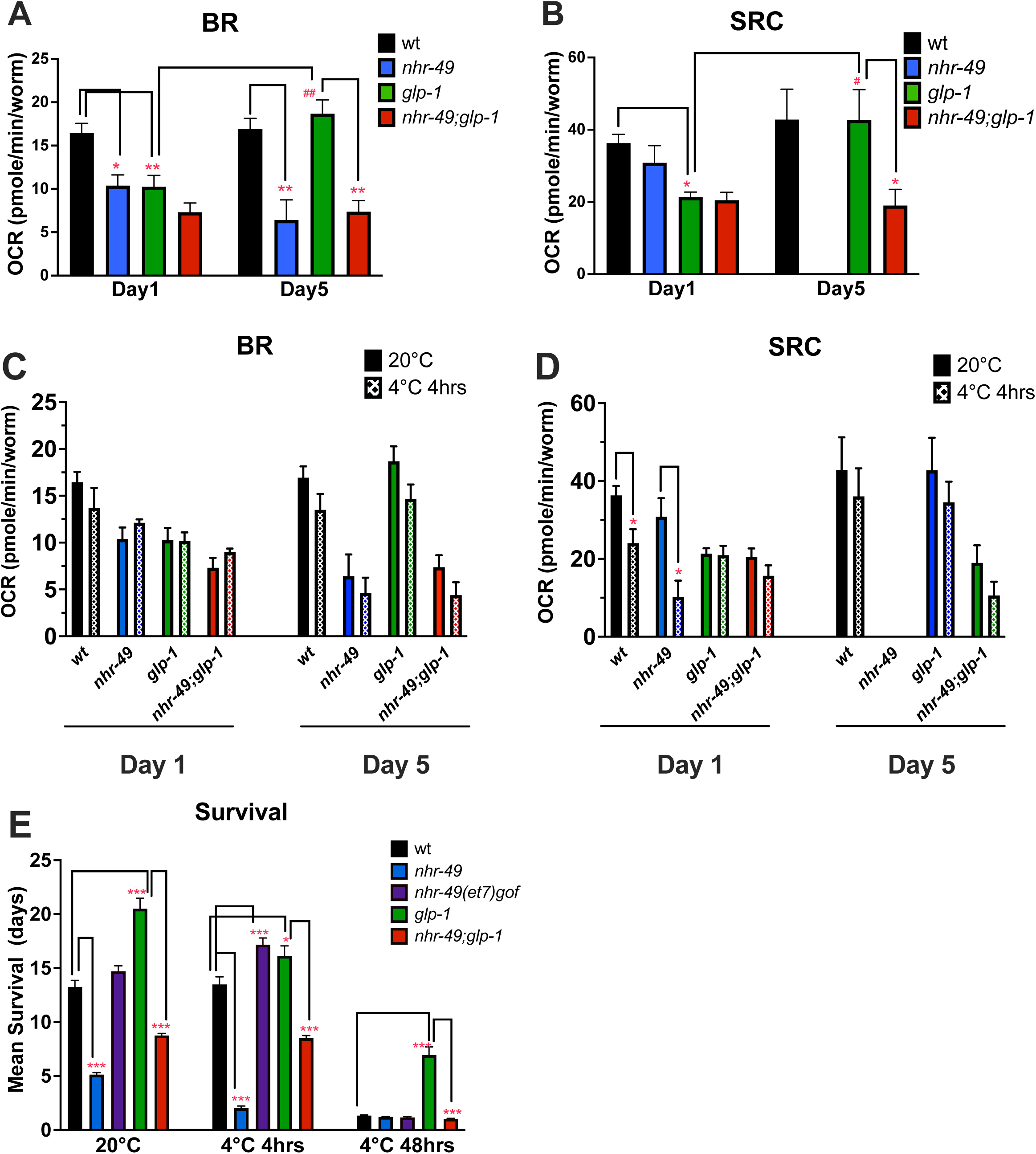
NHR-49 is required for maintaining mitochondrial function with age in germline-less animals. **A**-**D.** Basal respiration rate (BR; A, C) and spare respiration capacity (SRC; B, D) of wild-type animals (WT, black bars) and mutants of *nhr-49* (blue), *glp-1* (green) and *nhr-49;glp-1* (red) determined by measuring OCR measured using a Seahorsebio XF96 analyzer of Day 1 (left panels) and Day 5 adults, at normal temperature of 20^0^C (solid bars) and upon cold-shock exposure (4oC, 4h; hatched bars). Data shown as mean OCR ± standard error aggregated from independent trials (Day1 wt: n=5; Day1 *glp-1*: n=5; Day1 *nhr-49*: n=4; Day1 *nhr-49;glp-1*: n=4; Day5 wt: n=4; Day5 *glp-1*: n=4; Day5 *nhr-49*: n=3; Day5 *nhr-49;glp-1*: n=3). Statistical differences estimated by one-way ANOVA followed by Tukey’s multiple comparison in **A** and **B** and by Student’s T-test in **C** and **D**. **E:** Percentage of Day 1 adults of different stains surviving at control 20^0^C (left) or after exposure to 4oC for 4h (middle) or 48h (right). Survival analyses performed using Kaplan Meier survival tests and statistical significance determined by Mantel Cox method. Representative data from one of three trials. Data from additional trials in Table S1. Significance indicated as asterisks: * p<0.05, ** p<0.01, *** p<0.001, and **** p<0.0001.

### NHR-49/PPARα is essential for survival during acute and prolonged hypothermic stress

We assessed the impact of NHR-49 loss-of-function on organismal survival under hypothermic conditions. While *nhr-49* single mutants were ∼50% shorter-lived than WT worms at 20^0^C, they exhibited ∼70% lifespan reduction when exposed to acute cold shock (4^0^C, 4h) (Fig. 3E, Table S1). Alternatively, a strain carrying an *nhr-49* gain of function (*gof*) allele, *et7,* (Svensk et al., 2013) showed a modest survival improvement under these conditions (Fig. 3E, middle), suggesting that NHR-49/PPARα promotes hypothermic-stress resilience in wild-type animals. The survival in *nhr-49;glp-1* mutants was also significantly reduced by a 4hr cold exposure, though not as drastically as *nhr-49* single mutants (Figure 3E, Table S1). However, upon long-term exposure to cold (4°C for 48 h), *glp-1* mutants exhibited an extraordinary resistance and survived 50% higher than wild-type, and this extension was completely abrogated in *nhr-49;glp-1* (Fig. 3E, Table S1). Thus, NHR-49 is required for normal animals’ survival upon cold shock and is critical for the enhanced hypothermia resistance elicited by germline removal.

### DAF-16/FOXO3A does not impact mitochondrial function during hypothermia

Our previous studies showed that DAF-16/FOXO3A, a critical factor in the transcriptional network activated upon germline loss (Amrit and Ghazi, 2017; Ghazi, 2013), was necessary, in part, for the upregulation of *nhr-49* expression in *glp-1* mutants (Ratnappan *et al*., 2014). So, we examined the impact of a *daf-16* null mutation that completely abolishes *glp-1* mutants’ longevity under normal conditions (Arantes-Oliveira *et al*., 2002), on mitochondrial function. Surprisingly, neither the BR nor SRC was significantly reduced in *daf-16* or *daf-16*;*glp-1* adults, either at 20^0^C or upon exposure to 4^0^C for 4h in Day 1 (Fig. 4A, B) or Day 5 (Fig. 4C, D) adults. Therefore, DAF-16/FOXO3A does not appear to modulate the alternations in mitochondrial function induced by germline ablation under normal or acute hypothermic conditions.

**Figure 4.**
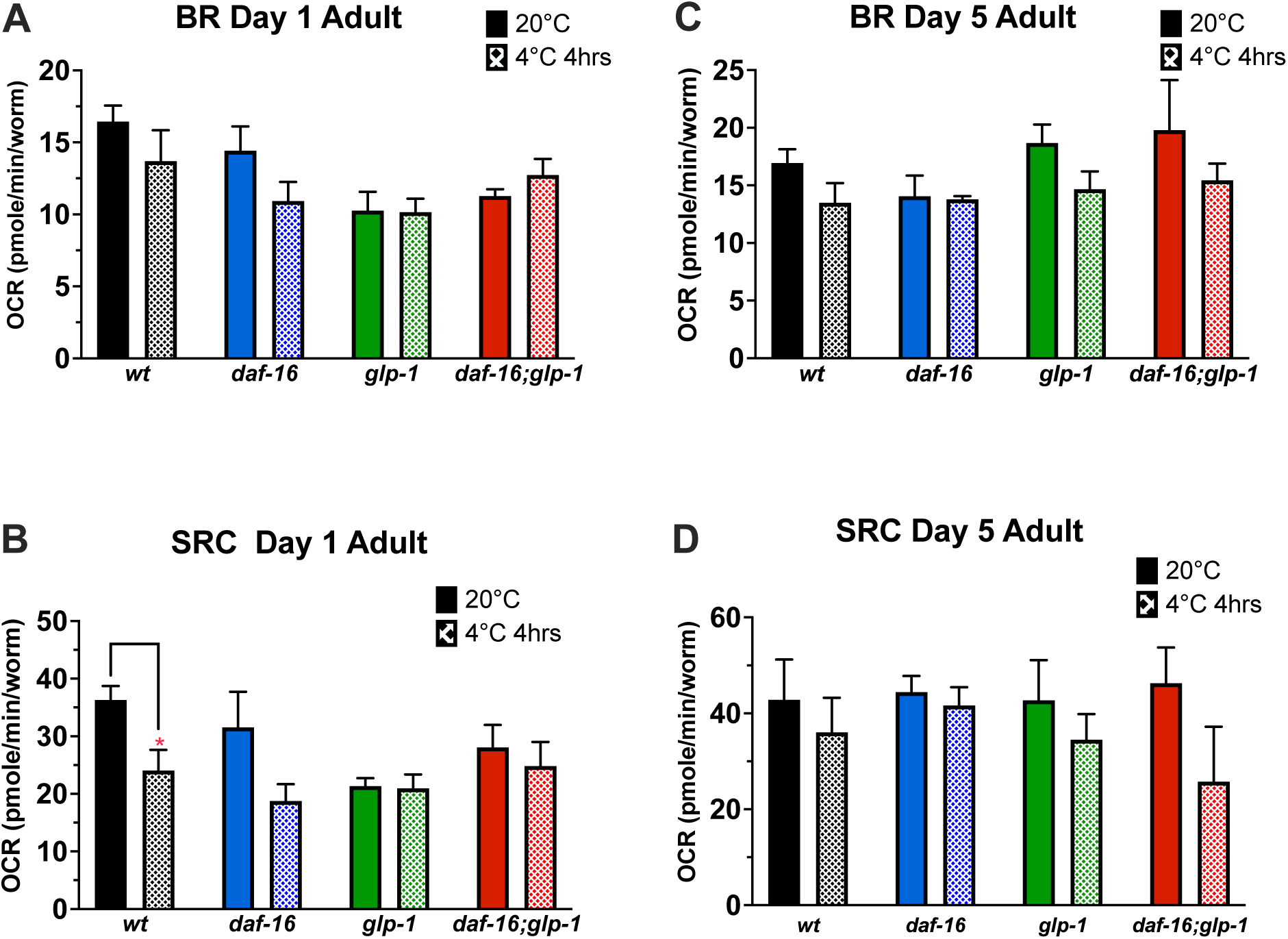
DAF-16 does not affect mitochondrial function in germline-less animals. Basal respiration rate (BR; A, C) and spare respiration capacity (SRC; B, D) of Day 1 (A, B) and Day 5 (B, D) adults of wild-type (WT, black bars) and mutants of *daf-16* (blue), *glp-1* (green) and *daf-16*;*glp-1* (red) strains determined by measuring OCR using a Seahorsebio XF96 analyzer at normal temperature of 20^0^C (solid bars) and upon cold-shock exposure (4oC, 4h; hatched bars). Data shown as mean OCR ± standard error aggregated from multiple 3-5 independent trials of each strain, age and condition (Day1 wt: n=5; Day1 *glp-1*: n=5; Day1 *daf-16*: n=3; Day1 *daf-16;glp-1*: n=3; Day5 wt: n=4; Day5 *glp-1*: n=4; Day5 *daf-16*: n=3; Day5 *daf-16;glp-1*: n=3). Statistical differences estimated by one-way ANOVA followed by Tukey’s multiple comparison and indicated as asterisks: * p<0.05, ** p<0.01, *** p<0.001, and **** p<0.0001.

## DISCUSSION

Traditional evolutionary theories of aging proposed an antagonistic relationship between fertility and longevity (Westendorp and Kirkwood, 1998). However, substantial evidence now shows that reproductive signals can impact health and aging both positively and negatively (Capezzuoli *et al*., 2022; El Khoudary, 2020; Falick Michaeli et al., 2015; Hsin and Kenyon, 1999; Mason *et al*., 2022; Ryan et al., 2018). Here, we found that, contrary to predictions based on our work and that of other laboratories (Amrit *et al*., 2016; Antebi, 2013; Goudeau *et al*., 2011; Ratnappan *et al*., 2016; Steinbaugh *et al*., 2015), germline loss leads to fragmented, poorly networked mitochondria and reduced function in early adulthood, but these animals maintain mitochondrial integrity and function later in life and during hypothermic stress. This highlights that reproductive status influences somatic physiology in both the short- and long-term, and triggers benefits or detriments, as seen in humans, where pregnancy and childbirth can have varying effects on women’s health across decades.

In *C. elegans* and other organisms, mitochondrial fragmentation typically correlates with poor respiratory function (Weir *et al*., 2017). Thus, it was surprising that long-lived, *glp-1* mutants— healthier than normal animals in many aspects (Amrit and Ghazi, 2017; Antebi, 2013; Ghazi, 2013)— display fragmented mitochondria. While an earlier reports suggested *glp-1* RNAi animals showed more elongated mitochondria (Chaudhari and Kipreos, 2017), our functional data support our anatomical findings, showing reduced mitochondrial volume and activity. Notably, sterile glp-4 mutants also exhibit reduced respiration but do not live longer (Shen *et al*., 2024). Interestingly, despite *glp-1* mutants’ increased resistance against multiple stressors (Arantes-Oliveira *et al*., 2002; Libina et al., 2003), an intact germline has been found to be required for mitochondrial stress to activate the UPRmt, and germline mitochondria have been implicated in this inter-tissue communication (Charmpilas *et al*., 2024; Shen *et al*., 2024). Similar paradoxes about mitochondrial fission-fusion dynamics and functionality. For instance, overexpression of mitochondrial fusion or fission genes both extends worm lifespan and increases stress resistance but, in both instances, despite fusion enhancement increasing respiration and ATP levels and fission elevation reducing respiration (Traa et al., 2024). Moreover, mitochondrial fragmentation is increased in both interventions. In fact, disruption of mitochondrial fission can also promote lifespan (Grohm et al., 2012).

Mitochondrial appearance and function do not necessarily represent or predict health or longevity, and local, tissue-specific changes likely shape whole-organism outcomes. Since we focused on muscle mitochondria alone, it is possible that other tissues show ostensibly ‘healthier’ appearance in *glp-1* mutants although our preliminary studies suggest intestinal mitochondria also show fragmented appearance (data not shown).

NHR-49/PPARα, a central regulator of mitochondrial fatty acid β-oxidation (Van Gilst *et al*., 2005a; Van Gilst *et al*., 2005b), is crucial for respiratory capacity and cold survival in germline-less animals. Unexpectedly though, it had age-specific requirements, since its loss had no perceptible impact on the BR of young *glp-1* mutants and was only essential for BR maintenance with age. Another surprising discovery was that DAF-16/FOXO was not involved here as our previous studied suggested that it acts upstream of NHR-49 in this pathway (Ratnappan *et al*., 2014) and has been implicated in age-related mitochondrial health in other longevity mutants (Wang et al., 2019). But, DAF-16/FOXO is also known to have contextual rather than universal impacts. GSC-depleted animals are able to induce somatic UPRmt in the absence of DAF-16/FOXO (Charmpilas *et al*., 2024) and its loss does not abrogate their enhanced heat-stress resistance either (Libina *et al*., 2003). It is possible that NHR-49/PPARα partners with one of the other NHRs or transcription factors activated upon GSC removal (Amrit and Ghazi, 2017; Ghazi, 2013). Overall, our findings highlight the intricate, dynamic relationship between reproductive status, mitochondrial metabolism, and longevity.

## EXPERIMENTAL PROCEDURES

### *C. elegans* Strains and Growth Conditions

All strains were maintained by standard techniques at 20°C on nematode growth medium (NGM) plates seeded with an *E. coli* strain OP50. Strains used in this study: wild type (N2), CF1903 *glp-1(e2144) III*, AGP12a *nhr-49(nr2041)*, AGP22 *nhr-49(nr2041)I;glp-1*, SJ4103 *zcIs14 [myo-3::GFP(mit)];N2*, AGP224 *zcIs14[myo-3::GFP(mit)];glp-1*, AGP 267 *zcIs14 [myo-3::GFP(mit)];nhr-49*, AGP268 *zcIs14 [myo-3::GFP(mit)];nhr-49;glp-1*, and AGP 269 *zcIs14 [myo-3::GFP(mit)];wt.* AGP224, AGP267, AGP268, and AGP269 were generated for this study by crossing SJ4103 and AGP22.

### Confocal Microscopy

Worms were mounted on glass slides with 2% agarose pads and anesthetized with 2 mM levamisole. Mitochondria in a body-wall muscle cell of a worm were imaged using a laser scanning confocal microscope (Confocal A1, Nikon) with 488 nm excitation and 525nm emission spectra. A muscle cell in anterior half of body was imaged. Images were analyzed using NIS-element AR software. Images were deconvoluted, and a single body wall muscle area in each image was defined as a region of interest (ROI) for z-stack slides. Green fluorescent objects within each ROI in the image were connected and the volume of 3D objects in one muscle cell was summed and averaged.

### Mitochondrial DNA Quantification

Mitochondrial DNA (mtDNA) from *C. elegans* was quantified using real-time quantitative PCR (qPCR). Synchronized populations of worms were harvested at the desired developmental stage, washed with M9 buffer, and lysed in a proteinase K solution at 65°C. Total DNA was extracted using a phenol-chloroform method, followed by ethanol precipitation. DNA concentration and purity were measured using a NanoDrop spectrophotometer. For qPCR, primers specific to the mitochondrial gene *mtDNA NADH dehydrogenase subunit 1* (mtND1) and a nuclear reference gene, such as *act-1* or *18S rRNA*, were used. Reactions were performed in a 20 µL volume with SYBR Green Master Mix on a real-time PCR machine. The relative mtDNA content was determined by calculating the ratio of mtND1 to the nuclear gene using the ΔΔCt method.

### Egg Synchronization and Sterilization

Worms were grown on NGM OP50 bacteria until day2 of adulthood. Day 2 Gravid animals were transferred to a fresh plate containing NGM OP50 and kept for 2h at 20°C for egg laying. After removing adult worms, the plates were kept at 25°C for 3 days to induce germline-stem cell loss in *glp-1* mutant worms. Day 1 adult worms were transferred to 20°C and maintained until being used for experiments.

### Oxygen Consumption Rates using XF96 Extracellular Flux Analyzer

Oxygen consumption rate (OCR) was measured on whole worms using the XF96 Extracellular Flux Analyzer (Agilent, Seahorse Bioscience) according to the manufacturer’s instruction with modification. Briefly, age-synchronized worms were washed with EPA water twice and transferred to an NGM petri dish without OP50. After 1hr, 5 worms were transferred to a XF96 plate well for 6∼8 technical replicates. OCR was measured by cycling of mix (2mins), wait (2mins) and measure (2mins) at room temperature. XF assay buffer consisted of 20 mM sodium pyruvate (#11360, Gibco) and EPA water (60mg MgSO4·7H2O, 60mg CaSO4·2H2O, and 4mg KCl.) The inhibitors of electron transport chain and oxidative phosphorylation system, carbonyl cyanide 4-(trifluoromethoxy) phenylhydrazone (FCCP, SigmaAldrich) (final concentration of 20 μM) and sodium azide (SigmaAldrich) (final concentration of 10 mM) were injected through a drug port after measuring 5 times of OCR. Each drug injection was performed in a separate experimental plate, in order to avoid a negative effect of FCCP on sodium azide. After adding drugs, OCR was measured after 8 times of cycling. OCR before adding drugs was measured as whole cellular respiration, OCR after adding FCCP was measured as maximal respiration, and OCR after adding sodium azide was measured as non-mitochondrial respiration. Mitochondrial basal respiration (BR) and spare respiratory capacity (SRC) were calculated by subtracting non-mitochondrial respiration from whole cellular respiration and by subtracting whole cellular respiration from maximal respiration respectively.

### Oxygen Consumption Rates using Clark-type Oxygen Electrode

For the OCR measurement using Clark-type oxygen electrode, mitochondria were isolated as previously described (Berry *et al*., 2023). OCR was normalized with protein concentration. Protein concentration was determined by Lowry assay.

Mitochondrial oxygen consumption was measured in a Clark-type oxygen electrode under several conditions. Succinate, pyruvate/malate, and palmitoyl carnitine were all used separately to measure the oxygen consumption dependent on complex II (succinate dehydrogenase), complex I (NADH ubiquinone oxidoreductase) and β-oxidation machinery, respectively. Gray arrows denote additions of substrate, ADP, Oligomycin, FCCP, and inhibitors. From the observed oxygen consumption rates, respiratory control ratios (RCR, State 3U/State4o) and reserve capacities (max rate-min rate, State3u-‘inhib’) were calculated and used as measures of mitochondrial quality and output, respectively. Data is represented as mean ± standard deviation of at least 5 independent mitochondrial preparations.

Adjusted P values are indicated on the graph

### Cold Stress Assays

Worms at the desired age were exposed to 4°C for 4h, recovered at 20°C for 2h, and utilized for OCR measurement and lifespan assay. For Day1 adults, after incubating at 25°C during development, worms were transferred to 20°C and incubated for 1h before exposure to 4°C.

## Acknowledgements

The authors are grateful to members of the Ghazi lab and the Pittsburgh ‘Wormclub’ members for valuable input throughout this study. Some strains were provided by the CGC, which is funded by NIH Office of Research Infrastructure Programs (P40 OD010440). This work was supported by a grant from the National Institutes of Health (R01AG051659) to AG and a Children’s Hospital of Pittsburgh Research Advisory Committee (RAC) postdoctoral fellowship to HH.

## SUPPLEMENTARY FIGURE AND TABLE LEGENDS

**Supplementary Figure 1:**
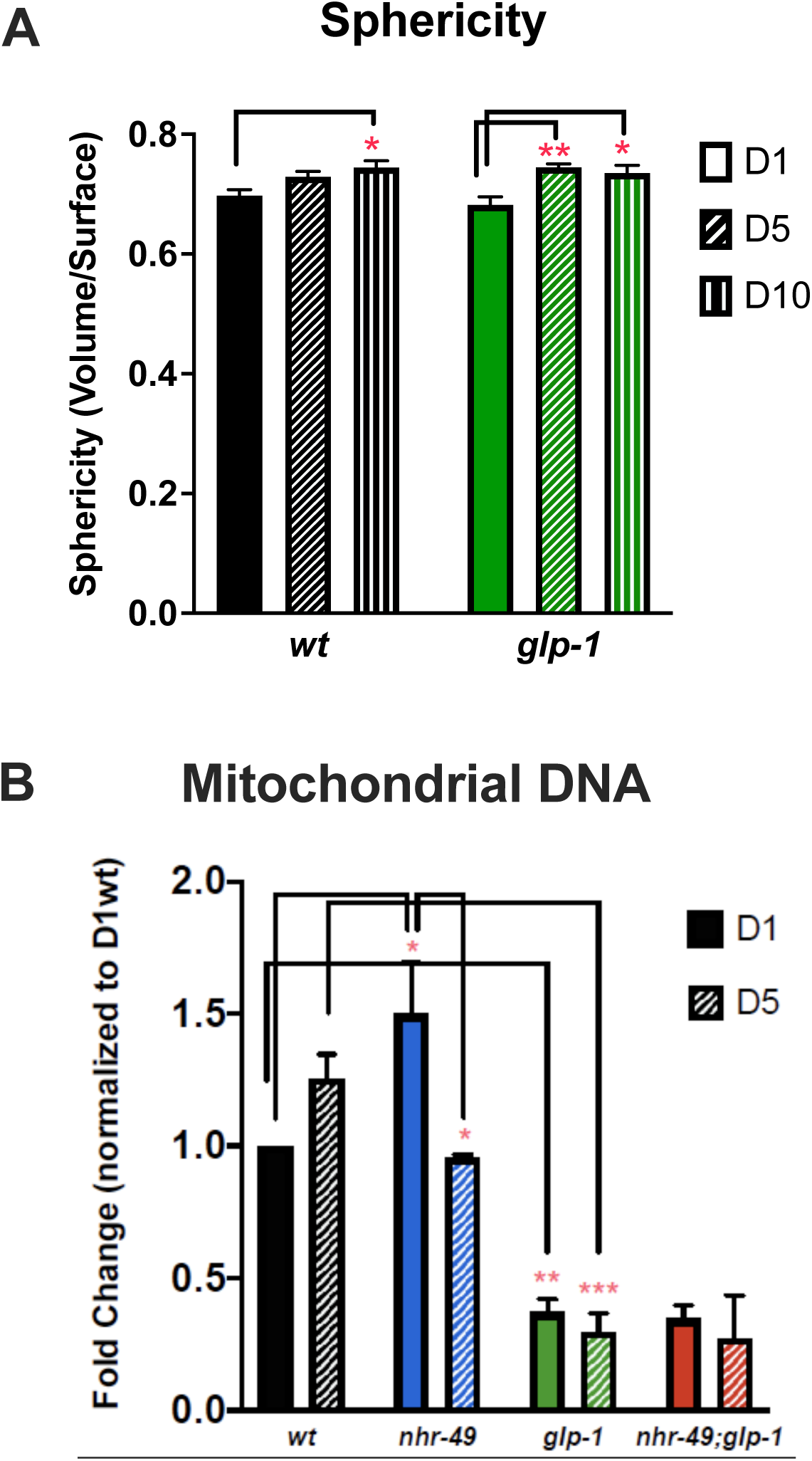
A: Sphericity of mitochondria measured from analysis of confocal images of the body wall muscle cell expressing Pmyo-3::mtGFP signals in wild-type (WT, black bars) and *glp-1* mutants (green bars) on Days 1(**B, top**), 5 (**B, middle**) and 10 (**B, bottom**) adults. **B:** Mitochondrial DNA quantification using Q-PCRs in Day 1(solid bars) or Day 5 (hatched bars) of different stains indicated. Statistical differences estimated by one-way ANOVA followed by Tukey’s multiple comparison and indicated as asterisks: * p<0.05, ** p<0.01, *** p<0.001, and **** p<0.0001. Data aggregated from 3-5 independent trials.

**Supplementary Figure 2:**
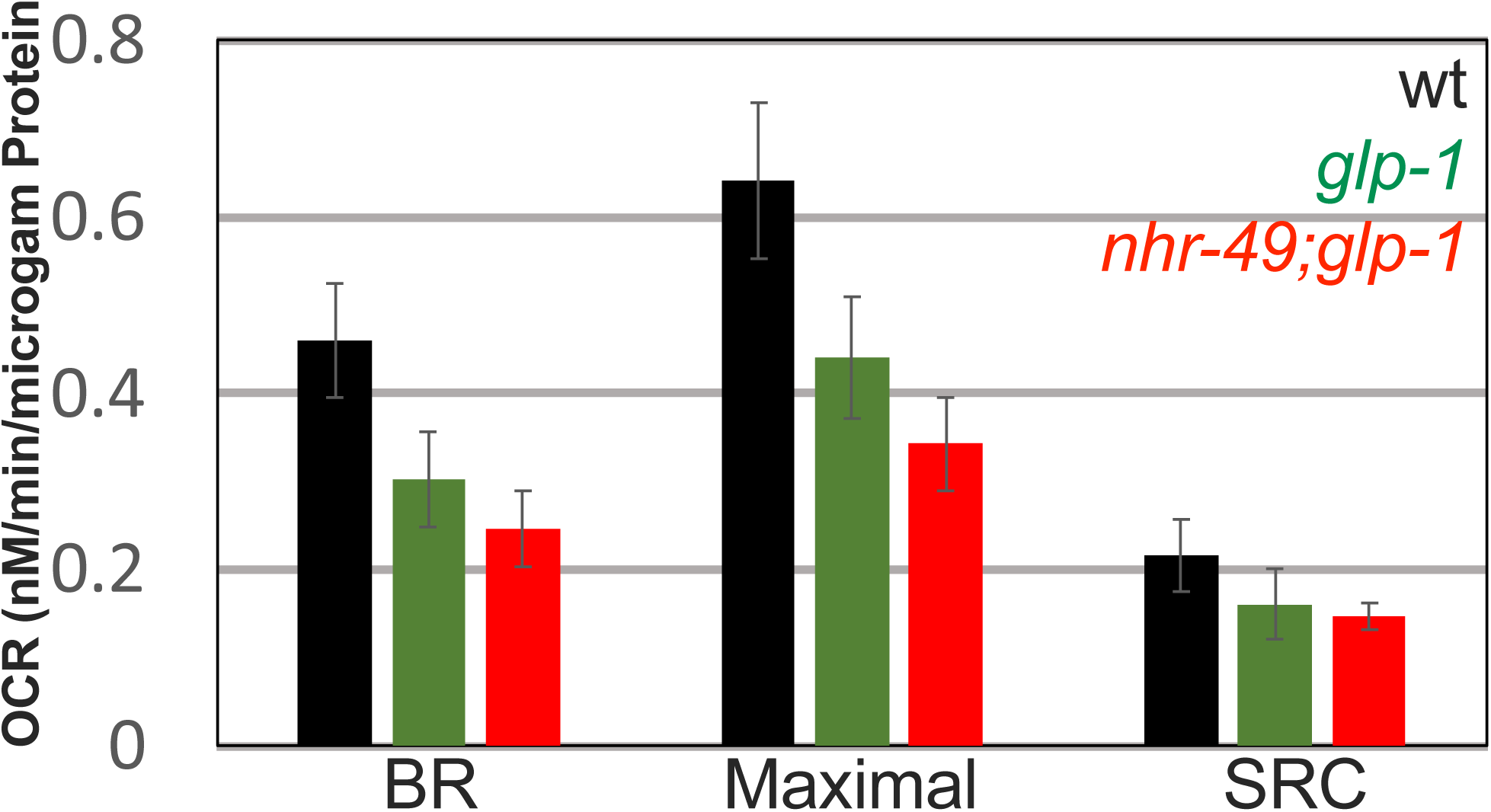
Basal Respiration (BR), Maximal Respiration (Maximal) and Spare respiratory capacity (SRC) of mitochondria isolated from Day1 adult worms of indicated strains using a Clark-type oxygen electrode. OCR was measured as (nmole/min/µg of proteins), with isolated mitochondria, for maximal respiration with FCCP (160µM) and non-mitochondrial respiration with sodium azide (20mM.). Columns indicate OCR average ± standard errors (wt: n=4, *glp-1*: n=4, and *nhr-49;glp-1*: n=5.)

**Table S1:**
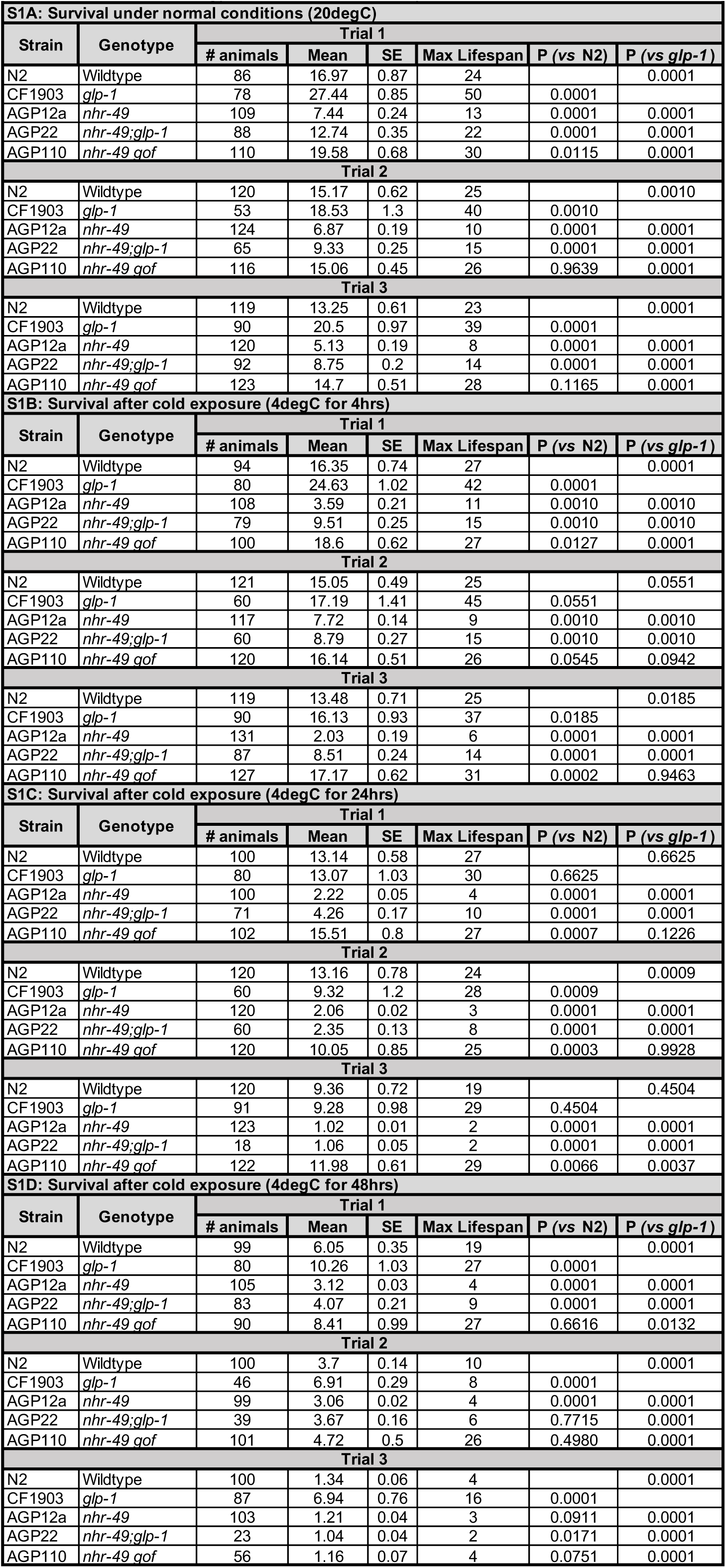
Survival of wildtype and mutant strains indicated at normal temperature (20^0^C; S1A) or upon exposure to 4oC for 4h (S1B), 24h (S1C) and 48h (S1D). Survival analyses performed using Kaplan Meier survival tests and statistical significance determined by Mantel Cox method with Bonferronic multiple comparison correction.

